# Descriptive analysis in time and space of recorded data for Buruli Ulcer occurrence in Victoria over 22 years

**DOI:** 10.1101/413542

**Authors:** Michael S. Avumegah, Ee Laine Tay, Soren Alexandersen, Wojtek P. Michalski, Daniel P. O’Brien, Isabelle Jeanne, Eugene Athan

**Affiliations:** Deakin University, Geelong, Australia.; Geelong Centre for Emerging Infectious Diseases (GCEID), Geelong, Australia.; Department of Health and Human Services, Melbourne, Australia.; Department of Infectious Diseases, Barwon Health, Geelong, Australia.; CSIRO Australian Animal Health Laboratory (CSIRO AAHL), Geelong, Australia.; Department of Medicine, Royal Melbourne Hospital, The University of Melbourne, Melbourne, Australia.

## Abstract

**Background:** Buruli ulcer (BU) is a subcutaneous necrotic infection of the skin caused by *Mycobacterium ulcerans*. There has been increasing BU incidence in Victoria, Australia. The aim of this study to provide an epidemiological update of BU cases in Victoria to understand the pattern of distribution over time and space and attempt to identify local risk factors.

**Methods:** A comprehensive descriptive epidemiological analyses were performed on BU notification data from 1994 to 2016. In addition, retrospective temporal, spatial and spatio-temporal analyses were conducted to understand the distribution of cases. Quantum GIS was used to generate maps. Demographic, new housing settlements and historical rainfall data were analysed to assess their effects on BU incidence in Victoria.

**Findings:** There were a total of 902 patients notified from 1994-2016. The incidence rate was 0.8/100,000 persons in Victoria. Space and time analyses showed that the most likely disease cluster was the Bellarine and Mornington Peninsulas with incidence rate 50 times higher than the State of Victoria rate. Gender was not a risk factor, but age was, with increased susceptibility among the over 60 year old group. There was an unusual high risk in the 15-24 age group in Point Lonsdale. Correlation analyses indicated that increase in population and construction of new settlements might be some of the reasons contributing to the rise in cases in Victoria.

**Interpretation:** The findings agreed with published works in Australia of the increase in BU cases in Victoria. However, our findings also highlights the endemic nature of cases. The identified spatial disease clusters could be relevant for future environmental sampling studies or screening tests for *M. ulcerans* exposure.

**Author Summary:** Buruli ulcer (BU) has been reported in 33 countries, mainly from the Tropics and Sub-tropics. Tropical cases are mainly within the West African region. Australia is the only country outside Africa in the top six highest incidence countries for BU. The exact mode of transmission remains unclear. Disease cases are rising in Australia, especially in Victoria for reasons that remains unclear. We have provided a descriptive epidemiological analyses in space and time of 22 years of recorded data on BU cases in Victoria from 1994 to 2016. We have also discussed demographic and new settlement dynamics over the study period. There were a total of 902 PCR-confirmed BU cases from 245 suburbs. Five suburbs on the Bellarine and Mornington Peninsulas were identified as the most endemic locations in Victoria. Spatial analyses detected a wider disease cluster area on the Peninsulas. We propose environmental sampling for risk factors analyses should focus on the endemic regions and some secondary clusters.

## INTRODUCTION

Buruli ulcer (BU) is a skin infection caused by *Mycobacterium ulcerans*. The progression of disease is marked by destruction of skin and subcutaneous tissue (1,2). While BU is the most common name, it is also known as Bairnsdale ulcer (2), Daintree ulcer (3), Kumusi ulcer (4), Mossman ulcer or Searle ulcer (5) depending on the geographical region (6).

A geographically restricted disease, BU has been reported in 33 countries in the tropics and sub-tropics (7,8), with very few cases reported in temperate areas (9). In the tropics, the highest prevalence is in the West and Central African regions (10,11). The sub-tropical cases are mainly within Queensland in northern Australia (3). Cases in the temperate regions include a temperate region in southern Australia (i.e., Victoria) as well as Japan and China (12). Australia reported its first case in Bairnsdale (Victoria) in the late 1930s (13,14). In Australia, MacCallum *et al.,* were the first to describe and publish reported cases of a skin ulcer caused by *Mycobacterium* cultivable only on Löwenstein-Jensen medium (2,14). The annual incidence rate as at 2016 is less than 1 per 100,000 inhabitants in Australia (15). Victoria is the state with highest number of cases reported in Australia (15).

The exact mode of transmission remains unclear; however, outbreaks of BU have usually been reported from areas close to aquatic environments (6,16). Aquatic insects have repeatedly been found positive for the *M. ulcerans* DNA target sequence IS*2404* (6,17,18). The involvement of mosquitoes as vectors and possums as reservoir have been suggested in Australia (12,19,20).

Heavy rains and floodings causing landscape alteration have also been associated with BU incidence (6,21). Age and gender are regarded as non-environmental risk factors in some areas (6,22). In Australia, BU is prevalent in the population older than 60 years; this has been attributed to immunosenescence (23).

BU cases are decreasing globally as per recent review (24), but Victoria has seen an increase in cases with shifting geographical distribution of disease (from the Bellarine to the Mornington Peninsulas) (15). Further studies are required to better understand this trend in Victoria. The main objective of this study was to provide an epidemiological profile and to perform space-time statistical analyses of BU cases in Victoria to understand distribution patterns and identify possible local risk factors such as age, gender, annual rainfall, changes in population and construction of new settlements.

## METHODOLOGY

### Ethics statement

Low risk ethics approval was obtained from Research Ethics, Governance & Integrity (REGI) Unit at Barwon Health for the use of the Victoria Buruli ulcer database. Project reference number: BW HREC 14/114. We were not required to obtain consent as data we were collating and analysing had already been notified to Department of Health and Human Services of Victoria (DHHS) under the Public Health and Wellbeing Act 2008 and patients were not re-contacted. All patient data analysed were anonymized.

### Buruli ulcer notifications for Victoria

The study population included patients diagnosed with BU and notified to the DHHS from 1 January 1994 to 31 December 2016. BU became a notifiable condition in Victoria in 2004, before which notification was voluntary. The BU confirmed cases involve detection from clinically suspected cases of IS*2404* by polymerase chain reaction (PCR) or isolation and culture of the organism from specimens performed by the *Mycobacterium* Reference Laboratory (MRL), Victoria. Data were extracted from a centralized notifiable disease database and variables used in the analysis included: BU notification date (stratified by month and year), 5 year age group, gender, suburb of residence and meshblock address at the time of notification, “exposure to which endemic area”, and “type of contact with endemic area (*Residents* or *Visitors*)”. Meshblocks are the smallest geographic regions in the Australian Statistical Geography Standard (ASGS), for which census data are available and preserve privacy. For purposes of interpretation, a “*Resident*” as referred to in this study is a BU patient with recorded exposure to *M. ulcerans* infection at their place of residence. There are two types of *Residents*: *Residents* living in a known endemic area and *Residents* who do not live in a known endemic area and never visited one. A “*Visitor*” is a BU patient who does not live in a known endemic area and have their exposure recorded outside their place of residence when visiting a known endemic area.

### Dataset for geographic information system

Geographic information system (GIS) map layers were acquired from the Government of Victoria website (**https://www.data.vic.gov.au/**) 25) and demographic data from Australian Bureau of Statistics (ABS) Data sourced included:

i. Regional boundaries (Australia and Victoria map layers)
ii. Meshblocks in Victoria (2011)
iii. Demographic data (2016 census data)
iv. New settlements information (2006-2016 census data)

Historical population and housing census data for Victoria from 1994 to 2016 has changed significantly. There has been the addition of new settlements and re-demarcation of meshblocks since 1994. Therefore, to provide a general epidemiological update, population census data used for all analyses were from the 2016 census data.

### Rainfall dataset and analyses

Since BU incidence has been linked to seasonal variations (6,21,27,28), historical rainfall data from 1994- 2016 were sourced from the Bureau of Meteorology website (29). Rainfall data for Rye was collected from the Rosebud country club station (#086213). Ocean Grove weather station (#087178) was also the source of Point Lonsdale and Queenscliff rainfall data. Barwon Heads rainfall data was collected from the Barwon Heads Golf Club station (#087135). The rainfall data quality were highly variable with 11 missing data points (1994, 1995, 1996, 1999, 2000, 2002, 2003, 2004, 2006, 2007 and 2009) from Ocean Grove weather station from 1994 to 2016. There were eight missing data points (1994 to 2001) from Barwon Heads Golf Club station and one missing data point (2003) from Rosebud country club station. To assess rainfall pattern and BU incidence at the local level, the mean annual rainfall from the three weather stations were cumulated into three groups namely 1994-2006, 2007-2011 and 2012-2016.

This grouping was done for ease of comparison with changes in the addition of new settlements from the ABS website from 2006, 2011 and 2016 housing and population census data at the local level.

### Statistical analysis and epidemiological calculations

GraphPad Prism (Version 7.03) was used to perform univariable analysis. The analyses included stratification by gender for age-group and temporal trend for entire Victoria as well as for the most endemic suburbs. Spearman correlation was performed for month of notification for BU cases stratified for *Residents* and the total number of places of residence as well as *Visitors* and total number of places of residence across Victoria. Nonparametric analyses was done using Kruskal–Wallis and Mann–Whitney tests on the variables at statistical significance at p-value of 0.05.

We also identified most common exposure sites and assessed demographic, physical environment and rainfall pattern. Epidemiological calculations for the most common exposure sites included: number of cases (n) and percentage (%) of positive cases in the population (pop), cumulative incidence rate for a time period, and relative risk. Cumulative incidence is the proportion of new cases occurring during a specified time period per 1,000 inhabitants. Relative risk is the ratio of percentage of positive cases among the exposed (to a risk factor) on percentage of positive cases among the non-exposed (**Table 1**). Relative risk and cumulative incidence were calculated for each endemic suburb and other detected disease clusters compared to the total population at risk in Victoria. All calculations were as described in Basic epidemiology (2^nd^ edition, World Health Organisation 2006) (30).

Mean annual cumulative incidence (Ci) = n ÷ (years under review) x 1,000 /population.

**Table 1:**
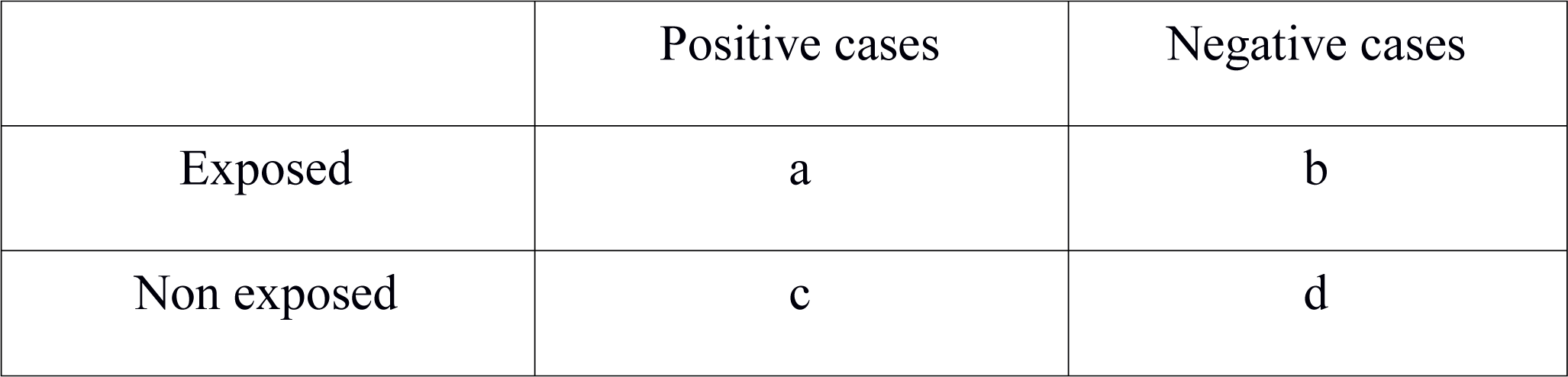
Relative risk contingency table.

Relative risk = a/(a+b)/c/(c+d)

We also assessed the correlation between BU incidence and changes in demographic, physical environment and rainfall pattern in the most common exposure sites.

### GIS and data importation

BU notification data containing the list of variables described above were imported into the GIS software by the meshblock unique identifier matching the Victoria meshblock layer. Extraction of the co-ordinates, latitude and longitude (WGS84 Geodesic reference system) of each polygon centroid from the two following layers: all meshblocks of Victoria and Victorian Buruli cases. Quantum GIS (qGIS) software (version 2.18.7, Las Palmas) was used.

### Organization of files for SatScan

To conduct the spatial, temporal, and space-time analyses, latitudes and longitudes were extracted for each meshblock centroid point using qGIS from a polygon map of Victoria. All BU cases locations were geo-coded and matched to the meshblock points and administration.

### SatScan analysis parameters

The files created as input files for SatScan software were:

**1. Case file** containing BU cases with meshblock location identification (Loc. id) and time of diagnosis in months and years.

**2. Population file** containing Loc. id with number of inhabitants

**3. Co-ordinates files** containing Loc. id with latitudes and longitudes

SatScan was programmed to report maximum cluster size of ≥ 25% of the population at risk. Spatial output was set to report hierarchical clusters with no geographic overlaps. Monte Carlo replications were set at 999.

### Spatio-temporal analysis

A retrospective purely temporal, purely spatial and space and time analysis under the Poisson probability model to detect significant clusters by exhaustively scanning over space, time and both space-time. Monte Carlo replication was performed using SatScan software (version 9.4.4). With a null hypothesis of no outbreak, the software algorithm searches the input data files to discover areas where cases exceed expectation and report a p-value and relative risk. In conjunction with qGIS, maps were developed to represent cases in time (temporal trend), in space (spatial analysis) and in space and time (spatio-temporal trend) (31).

## RESULTS

### Distribution of Buruli ulcer cases by residence in Victoria-1994-2016

There were 902 confirmed BU cases notified from 1994 to 2016 with residential addresses from 245 suburbs in Victoria (**Fig 1**).

**Fig 1:**
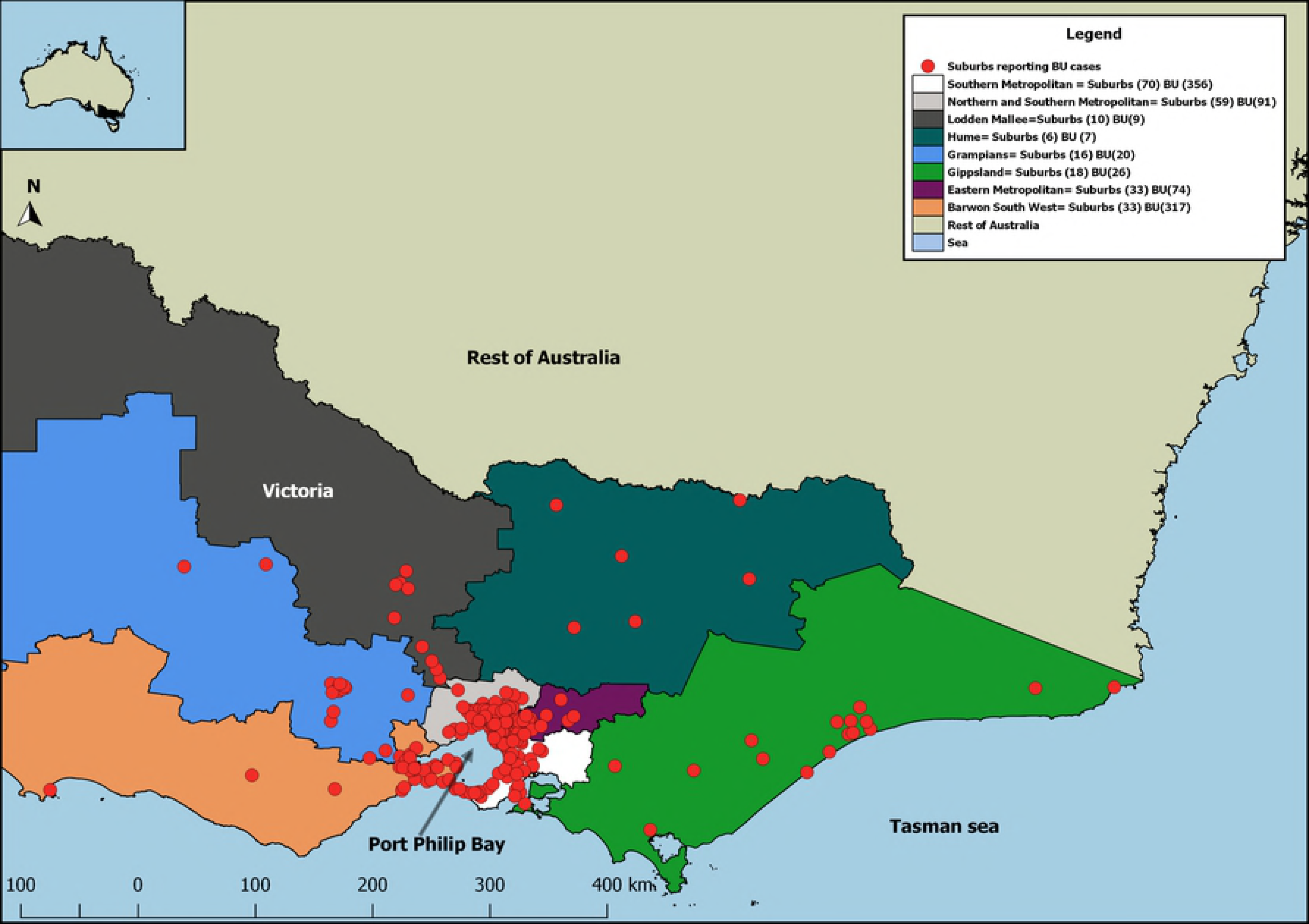
Distribution of confirmed Buruli ulcer cases within Victorian Government regions in Victoria from 1994 to 2016. The coloured regions shown in the map represent the Government of Victoria regions in 2011.

The highest concentration of reported cases lived in the Southern metropolitan region of Melbourne, Victoria, with 356 cases (39%). This was closely followed by Barwon South West region with 317 cases (35%). Northern/Southern and Eastern metropolitan had 91 (10%) and 74 (8%) respectively. Gippsland, Grampians, Loddon-Mallee and Hume regions reported < 30 cases (**Fig 1**). Of these cases, 523 (58%) had their exposure location recorded as their suburb of residence, 335 (37%) had their exposure location recorded outside their suburb of residence and 44 (5%) had no recorded exposure site. More males (509; 56%) reported cases compared to females (392; 44%). The three age groups with most cases were 60-64, 65-69 and 70-74 years recording 83 (9%), 78 (9%) and 72 (8%) cases, respectively. The remaining age groups had < 70 cases each (**Fig 2A**). There were no significant differences in age, when stratified by gender (p = 0.13). The same was observed for month of notification (p = 0.20) and year of case report (p = 0.61). Month of notification of BU cases correlate with *Residents* and the total number of different suburbs they resided as well as *Visitors* and the total number of different suburbs they resided across Victoria (**Fig 2D**).

**Fig 2:**
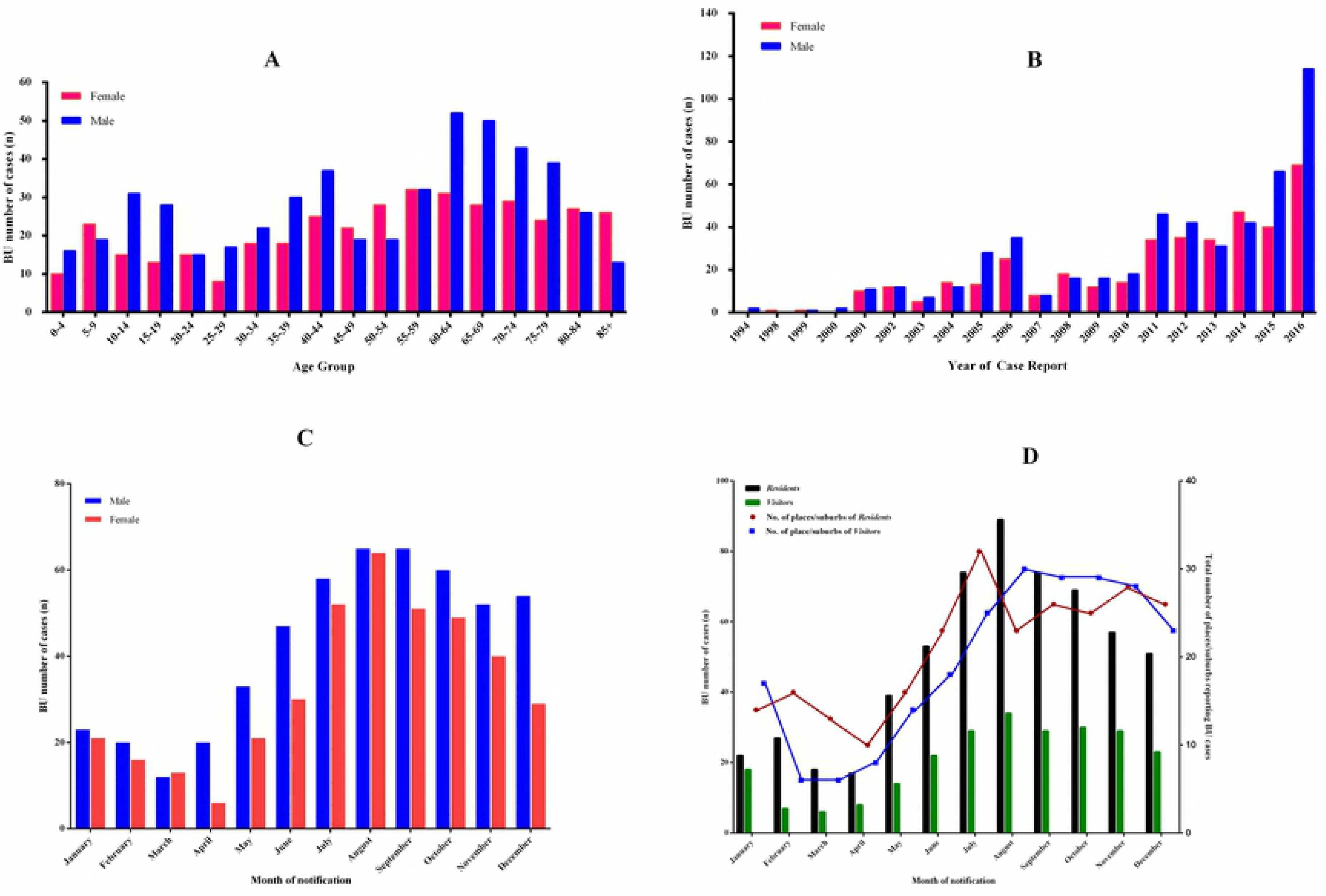
A: Age group stratified for gender. B: Year of cases report stratified for gender. C: Month of notification stratified for gender. D: Month of notification for BU cases stratified for *Residents* and the total number of places of residence as well as *Visitors* and the total number of places of residence across Victoria.

### Detected purely spatial and space-time clusters

The purely spatial and space-time analysis showed an average annual reported incidence of BU cases per 100,000 people of 0.8 from 1994 to 2016 in Victoria. The space-time analysis reported five spatial disease clusters (1, 3, 4, 5 and 6) and one temporal cluster (cluster 2). The purely spatial analysis identified 11 clusters. Four of the space-time clusters (1, 3, 4 and 5), as well as the purely spatial clusters (1, 2, 3 and 4) were reported to be significant clusters (p < 0.05, **Fig 3**). Both the purely spatial and space-time analysis had the most likely disease cluster (cluster 1), also known as the primary/candidate cluster geographically located on the Bellarine and Mornington Peninsulas (**Fig 3**). The annual incidence for cluster 1 (**Fig 4**) of reported cases/100,000 persons for the space-time and purely spatial was 40 (2001-2016) and 28, respectively. Temporal analysis detected significant rise in BU cases from 2011 to 2016 (red lines) (**Fig 4**).

**Fig 3:**
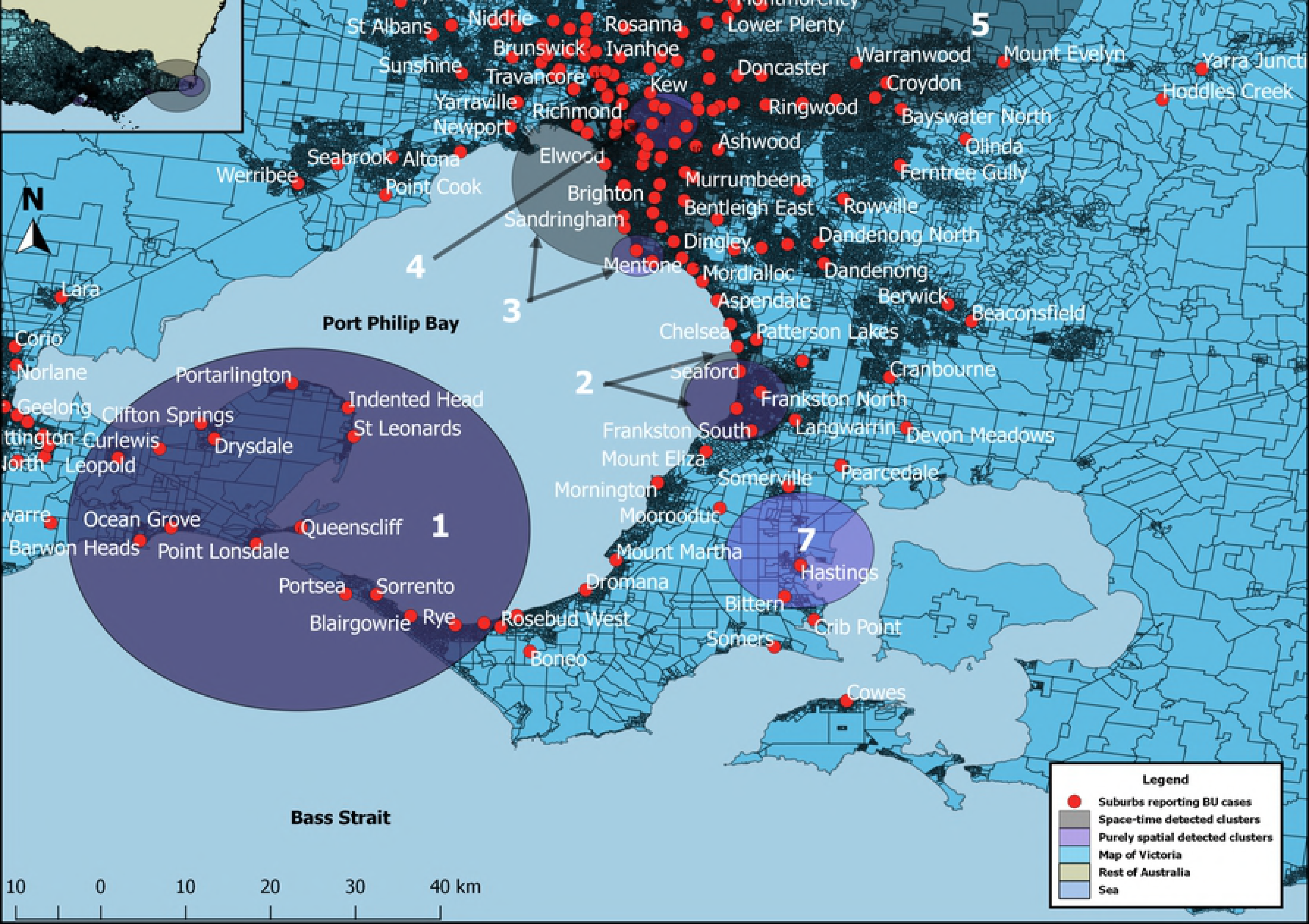
Detected patients’ clusters of purely spatial and space-time analyses. Both primary clusters of space-time and spatial analyses labelled “1” (Indigo circular window) are located on the Bellarine and Mornington Peninsulas. The remaining circular windows (Violet circular window: purely spatial clusters and Black circular windows: space-time clusters) seen on the map are secondary clusters. The numbers (1, 2, 3, 4, 5, 6 and 7) represent the order of cluster reporting. Some secondary clusters could not be shown on map due to small size.

**Fig 4:**
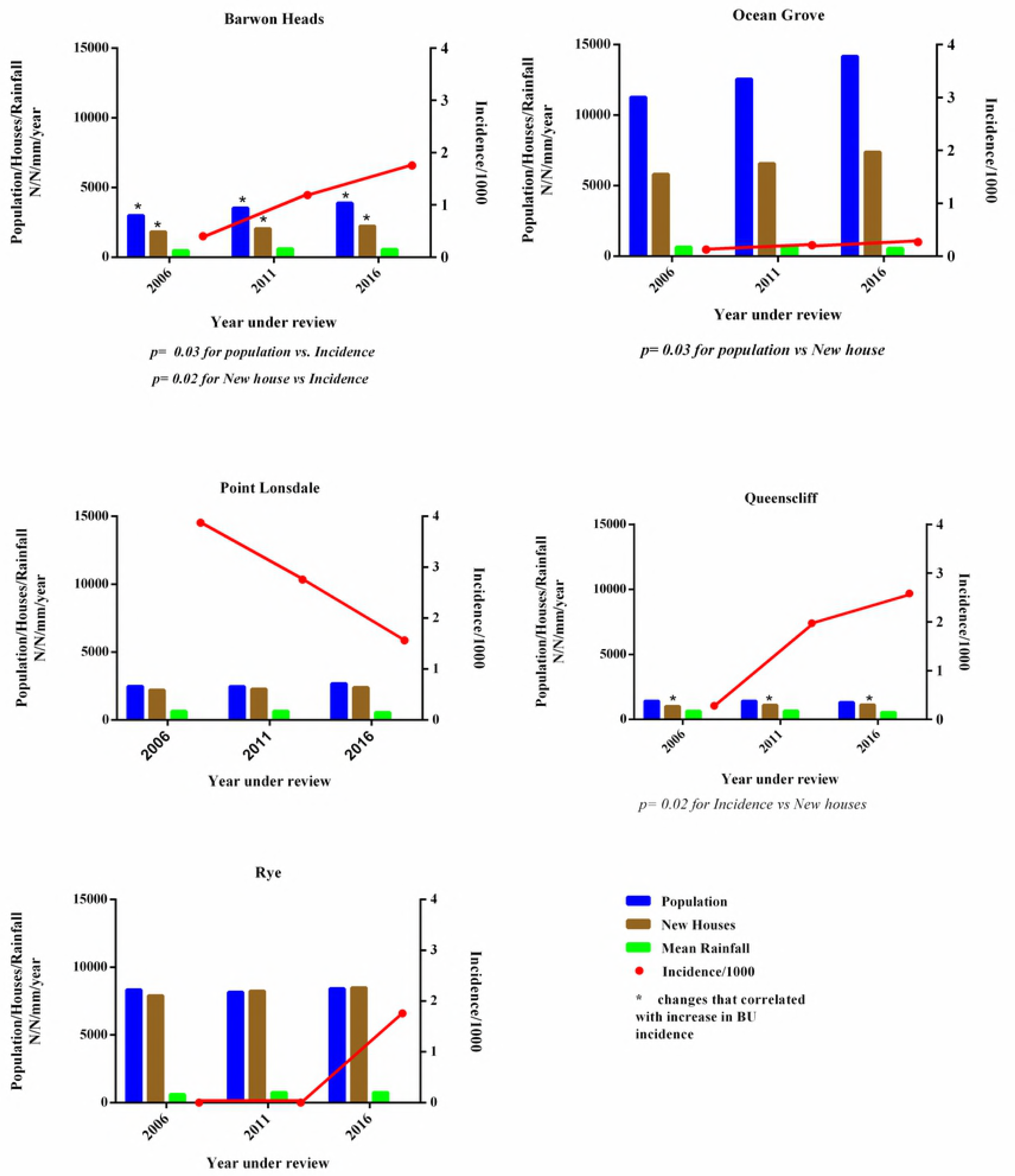
Purely temporal trend of BU cases from 1994 to 2016. The lines represent observed cases across time: Blue line shows observed cases only. Red line indicates observed cases that showed significant temporal clustering across time (also highlighted in red). Dark green line indicates the baseline expected cases based on the total population used in the analysis (default calculation by SatScan) and (iv) Light green line is the ratio of observed ÷ expected BU cases.

### Top five residence and exposure locations

The top five suburbs for the place of residence and exposure location of most reported BU cases were located on the Bellarine and Mornington Peninsulas from 2002-2016 period (**Fig 5**).

**Fig 5:**
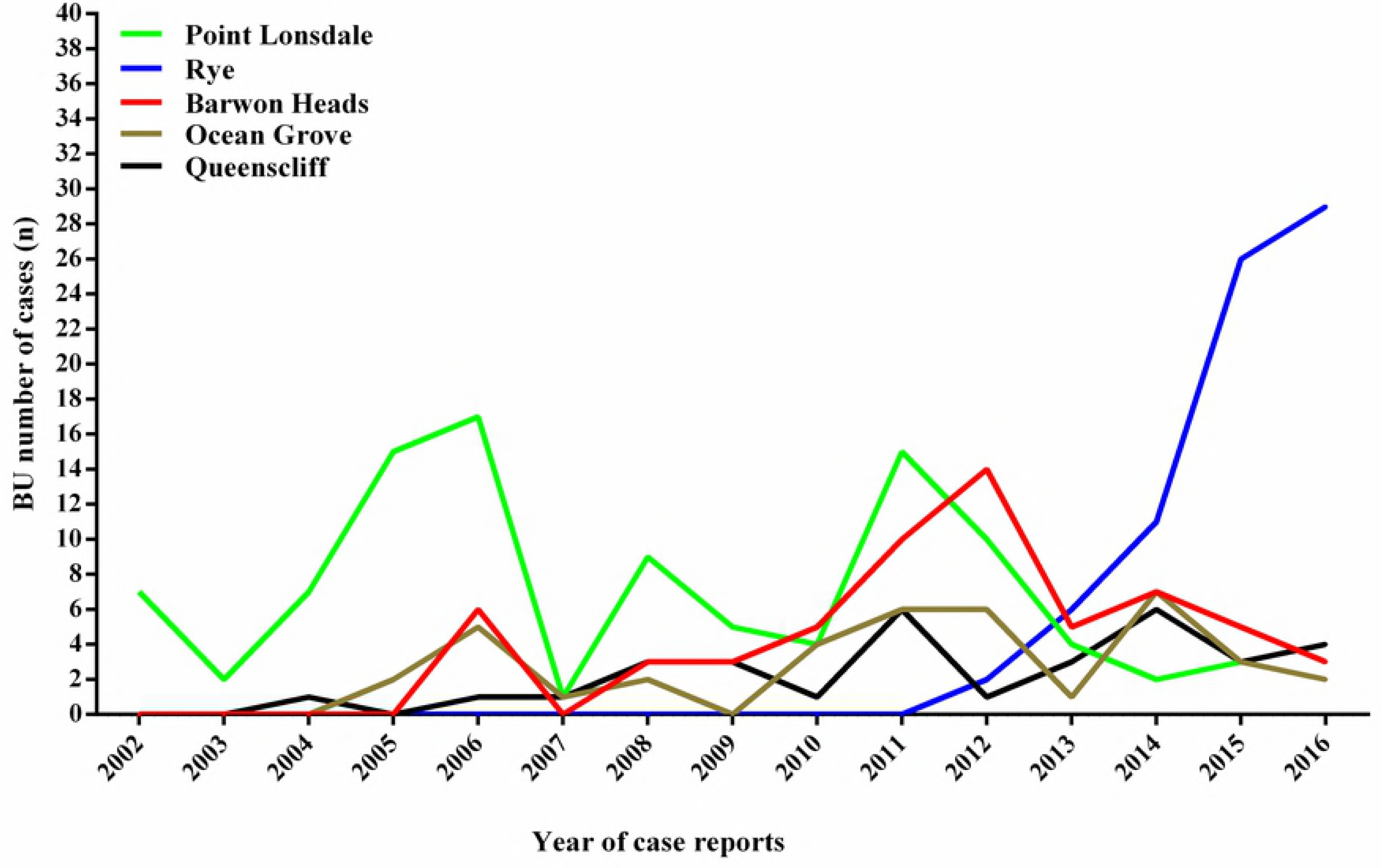
Number of BU cases in the top 5 residence (5A) and most visited/attributed exposure (5B) areas from 1994 to 2016. Fig 5B, shows the most visited or attributed *M. ulcerans* exposure areas.

The suburb with most BU cases in *Residents* was Point Lonsdale with 103 (11%) cases, followed by Rye with 74 (8%), Barwon Heads with 61 (7%) Ocean Grove with 39 (4%) and Queenscliff with 33 (4%) cases. All the patients in these suburbs had their exposure location recorded as their place of residence in the Bellarine or Mornington Peninsulas (**Fig 5A**). **Fig 5B** shows the top five suburbs reported to be the place of probable exposure by patients (*Visitors*) not living in the area. Point Lonsdale was the suburb with the highest number of perceived exposure cases in *Visitors* with 89 (10%) cases followed by Barwon Heads with 42 (5%) cases. Rye, Ocean Grove and Queenscliff followed with 32 (4%), 24 (3%) and 24 (3%) cases, respectively.

### Temporal trend in the top five *Resident* and most visited areas

The temporal trend of *Residents* and *Visitors* locations were compared as well as years of case report for each of the five suburbs (**Fig 6**). Since 2011, Point Lonsdale has been reporting a decrease in cases whereas Rye, has been reporting increasing cases since 2012 (**Fig 6**). The seasonal trend of BU reporting was generally similar in both *Residents* and *Visitors* (**Fig 2D**).

**Fig 5:**
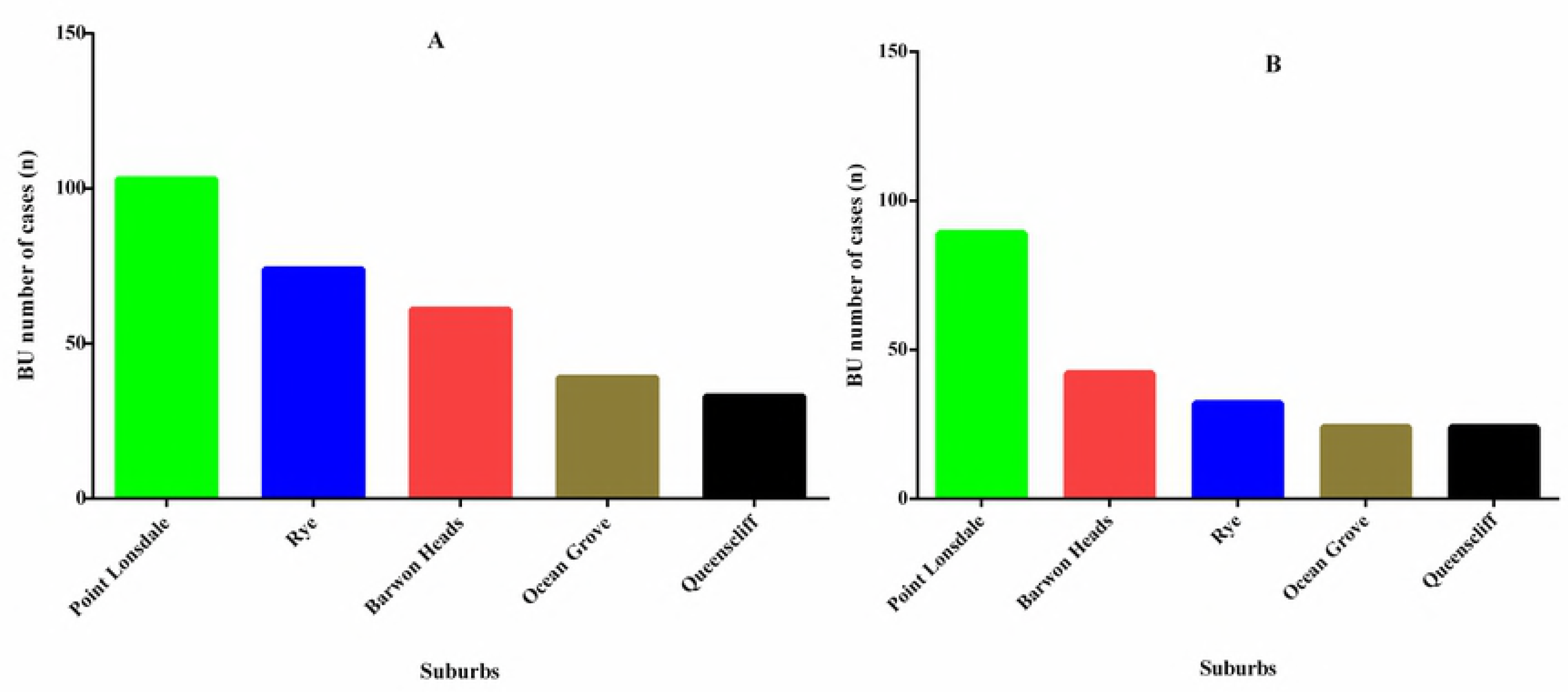
Five top suburbs of BU cases from 2002 to 2016. Number of diagnosed cases among *Residents* and *Visitors* in the top five residence and perceived exposure areas in each month from 1994-2016. Cases in Point Lonsdale (green) has shown a decrease since 2011, while Rye (blue), which started reporting cases in 2012 has shown a steady increase. BU cases in Barwon Heads (red), Ocean Grove (brown) and Queenscliff (black) have been dynamic since the onset of case reporting.

### Age and gender of *Residents* and *Visitors* in the top five suburbs

From the 2016 census data, the most populated suburb in the top 5 *Residents’* locations was Ocean Grove (14, 165), followed by Rye (8, 416) (**Table 2**). Rye had almost twice as many males to females among both *Residents* and *Visitors* reporting BU. The proportion of males to females with BU cases among Resident or *Visitors* was similar within each of the other four suburbs (**Table 2**). There was no statistically significant difference in gender among *Residents* (p= 0.44) as well as *Visitors* (p= 0.52, **Table 2**). There was a significant difference in age group of *Residents* (**Table 2**, p= 0.04) and *Visitors* (**Table 2**, p= 0.00).

**Table 2:**
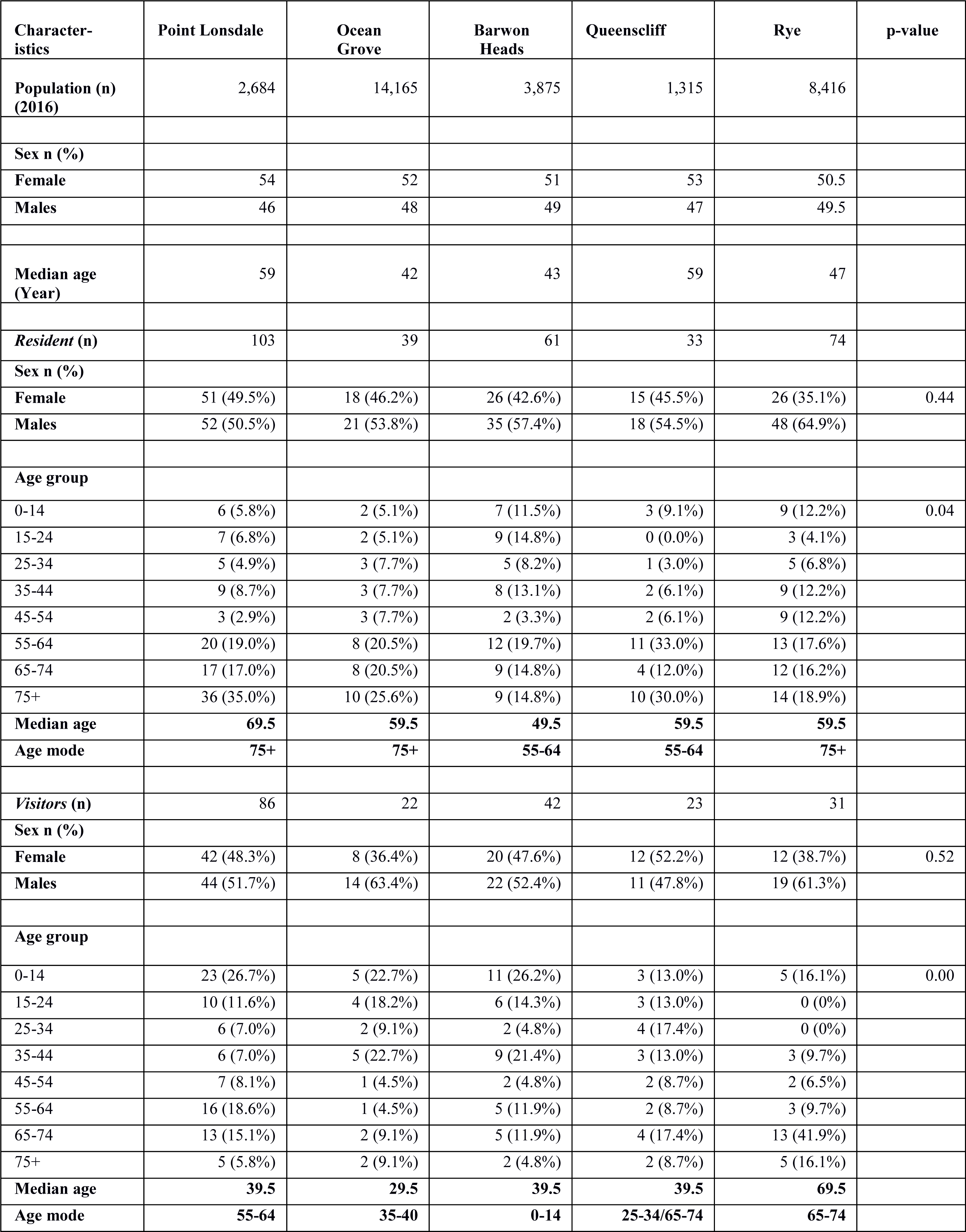
Age and gender of *Residents* and *Visitors* in the top five suburbs.

### Changes in demographics and physical environment of the top 5 exposure sites

Within a ten year time period (i.e., 2006 to 2016), Barwon Heads and Ocean Grove have had the greatest population increases of 881 (21%) and 2,891 (20%) respectively (**Fig 7**). Over the same time, the population at Point Lonsdale increased by 207 (8%) and Rye 79 (1%). There has been an 8% decrease in Queenscliff population in the same time period. Ocean Grove had the greatest number of new house constructions at 1,574 (21%). Rye, which had only 1% increase in population had a 590 (7%) new constructions. There was a significant correlation between the incidence rate of BU cases and the increase in population and new housing constructions in Barwon Heads (p = 0.02). There was also a significant correlation between incidence rate and new housing constructions in Queenscliff (**Fig 7**, p = 0.02). The remaining suburbs did not show any significant relationship between incidence rate of cases, population change and housing constructions. The rainfall pattern did not significantly affect incidence rate in top five areas at the local level (**Fig 7**).

**Fig 6:**
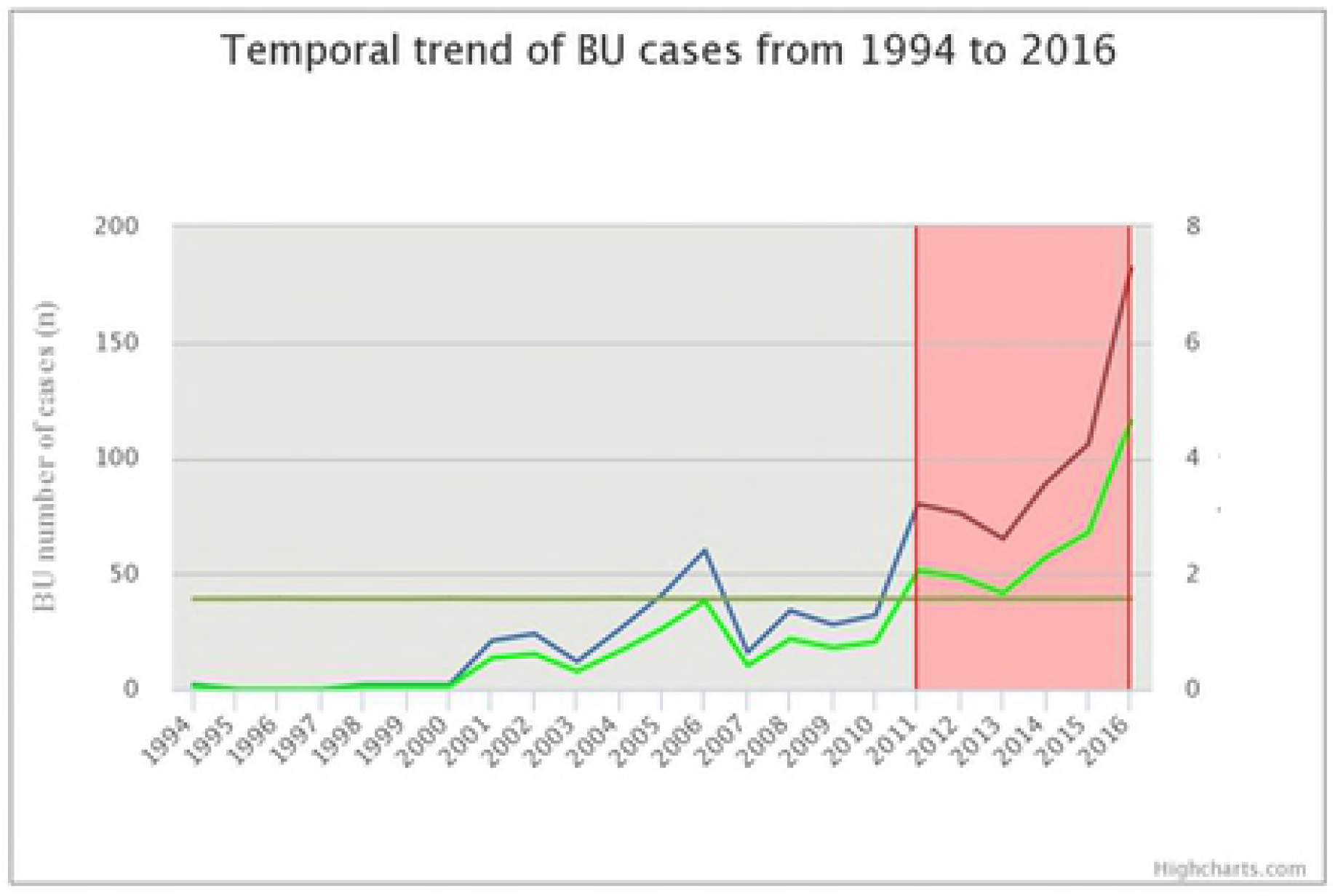
Incidence of BU cases with changes in population, new house constructions and rainfall data from 2006 to 2016. The year under review were considered for “2001 to 2006”, “2007 to 2011” and “2012 to 2016”. The blue, brown and green bars represent population (N), new house constructions (N) and rainfall historical data (mm/year), respectively. The incidence rate of BU cases is shown in red dots and lines. Correlation analyses was performed for each variable to understand how they relate to incidence rate. P-values (*) were reported where the relationship was significant (p < 0.05).

### Relative risk of living in the top 5 or disease cluster areas

The relative risks recorded for BU in *Residents* in each of the top five patients’ locations and disease cluster areas compared to the rest of Victoria were high. The risk of having been diagnosed with a Buruli ulcer was 274 (range: 224 - 335) times higher in Point Lonsdale compared to the rest of Victoria. This risk in Queenscliff and Barwon Heads were 171 (122-241) and 111 (86-143), and for Ocean Grove and Rye, it was less than 100. Relative risk of those living in the top five areas (combined) was 101 (88 - 116).

The highest absolute risk of BU cases was 4 % in Point Lonsdale, 3 % in Queenscliff and 2 % in Barwon Heads. Ocean Grove and Rye had less than 1% each. The highest mean annual cumulative incidence of 2.6/1,000 persons was observed in Point Lonsdale. Queenscliff, Rye and Barwon Heads had 1.9, 1.8 and 1.4/1,000 persons respectively. The lowest cumulative incidence of 0.2/1000 persons was recorded in Ocean Grove (**Table 3**).

**Table 3:** Relative risk of living in the top 5 or disease cluster areas.

| Location or cluster | Resident Population (N) | Period in review | Total cases (n) | Mean cases number/ year | Absolute risk (%) | Mean annual incidence/ 1000 inhabitants | Relative risk (Ci 95%) |
| --- | --- | --- | --- | --- | --- | --- | --- |
| Barwon Heads | 3,875 | 2006-2016 | 61 | 5.5 | 2 | 1.4 | 111 (86 -143) |
| Ocean Grove | 14,165 | 2005-2016 | 39 | 3.3 | < 1 | 0.2 | 19 (14 -26) |
| Point Lonsdale | 2,684 | 2002-2016 | 103 | 6.9 | 4 | 2.6 | 274 (224 -335) |
| Queenscliff | 1,315 | 2004-2016 | 33 | 2.5 | 3 | 1.9 | 171 (122 -241) |
| Rye | 8,416 | 2012-2016 | 74 | 14.8 | 1 | 1.8 | 63 (50 -80) |
| Combined (5) | 30,455 | 2002-2016 | 310 | 20.7 | 1 | 0.7 | 101 (88 -116) |
| **Space-Time** |  |  |  |  |  |  |  |
| Cluster 1 | 61,723 | 2001-2016 | 399 | 26.6 | < 1 | 0.4 | 75 (66 -86) |
| Cluster 2 | - | 2011 2016 | 598 | 99.6 | - | - | - |
| Cluster 3 | 73,072 | 2005-2016 | 62 | 5.2 | < 1 | 0.1 | 6 (5 - 8 ) |
| Cluster 4 | 638,943 | 2016 | 50 | 50 | < 1 | 0.1 | 1 (0 - 1) |
| Cluster 5 | 18,033 | 2002-2010 | 14 | 1.6 | < 1 | 0.1 | 5 (3 - 9) |
| **Purely spatial** |  |  |  |  |  |  |  |
| Cluster 1 | 61,723 | 1994-2016 | 402 | 17.5 | 1 | 0.3 | 76 (67 -87 ) |
| Cluster 2 | 64,706 | 1994-2016 | 63 | 2.7 | < 1 | 0.0 | 7 (5 - 9) |
| Cluster 3 | 16,140 | 1994-2016 | 24 | 1.0 | < 1 | 0.1 | 10 (7 -15) |
| Cluster 4 | 101,677 | 1994-2016 | 53 | 2.3 | < 1 | 0.0 | 4 (3 - 5) |

The Space-time and purely spatial primary cluster, also known as the most likely cluster (Cluster 1, **Fig 3**) where observed cases are significantly higher than expected, which circumscribes areas on the Bellarine and Mornington Peninsulas had relative risk of 75 (66 - 86) and 76 (67 - 87) and mean annual cumulative incidence of 0.4 and 0.3/1,000 persons respectively. Relative risk of the remaining clusters was ≤ 10. Cumulative incidence recorded in cluster 2, 3 and 4 of the purely spatial analysis were less than 1 (0.0- 0.3/1,000 persons, **Table 3**). The same could be said for the remaining space-time clusters (3, 4 and 5, **Fig 3**) with 0.1/1,000 persons (**Table 3**).

### Absolute risk and incidence rate for age groups and gender within the top five *Residents’* locations

The highest absolute risk of 7% was found among the > 75 year group in Point Lonsdale (**Table 4**). Queenscliff had the next highest risk of 5% among the same age group. The next age group with the second highest absolute risk was among the 55-64 year group found in Point Lonsdale (5%) and Queenscliff (4%). The third highest absolute risk of 4% was found in Point Lonsdale among the 15-24 year group. The remaining absolute risk varied considerably among the rest of the suburbs (range: 0 to 4%) (**Table 4**). The absolute risk of 3% for all the suburbs combined was among the > 75 year group followed by 55-64 and 65-74 year groups with both having 1% each (**Table 5**). Absolute risks and incidence rate among gender was not statistically significant (**Table 6**, p = 0.81). Age was a risk factor in the five endemic areas (**Table 4**, p = 0.00).

**Table 4:** Buruli ulcer absolute risk (%) for age groups within each of the top five *Residents’* location using 2016 population census data.

| Age group | PL |  |  | OG |  |  | BH |  |  | QC |  |  | Rye |  |  |
| --- | --- | --- | --- | --- | --- | --- | --- | --- | --- | --- | --- | --- | --- | --- | --- |
|  | n | pop | % | n | pop | % | n | pop | % | n | pop | % | n | pop | % |
| 0-14 | 6 | 364 | 2% | 2 | 2,904 | <1% | 7 | 928 | 1% | 3 | 139 | 2% | 9 | 1,393 | 1% |
| 15-24 | 7 | 168 | 4% | 2 | 1,327 | <1% | 9 | 348 | 3% | 0 | 100 | 0% | 3 | 691 | < 1% |
| 25-34 | 5 | 138 | 4% | 3 | 1,358 | <1% | 5 | 237 | 2% | 1 | 58 | 2% | 5 | 775 | 1% |
| 35-44 | 9 | 233 | 4% | 3 | 1,990 | <1% | 5 | 562 | 1% | 1 | 77 | 1% | 5 | 1,023 | 1% |
| 45-54 | 3 | 282 | 1% | 3 | 1,844 | <1% | 2 | 523 | <1% | 2 | 167 | 1% | 9 | 1,055 | 1% |
| 55-64 | 20 | 395 | 5% | 8 | 2,011 | <1% | 12 | 566 | 2% | 11 | 250 | 4% | 13 | 1,218 | 1% |
| 65-74 | 17 | 542 | 3% | 8 | 1,680 | 1% | 9 | 450 | 2% | 4 | 299 | 1% | 12 | 1,325 | 1% |
| 75+ | 36 | 543 | 7% | 10 | 1,061 | 1% | 9 | 246 | 4% | 10 | 213 | 5% | 14 | 933 | 2% |
| **Total** | **103** | **2,665** |  | **39** | **14,175** |  | **58** | **3,860** |  | **32** | **1,303** |  | **70** | **8,413** |  |

n= number of BU cases for that age group
pop= total number of *Residents* with that age group in the suburb
% = absolute risk calculated for that age group in the suburb (n/pop) x 100.
PL= Point Lonsdale
OG= Ocean Grove
BH= Barwon Heads
QC= Queenscliff
Age group is a risk factor in the endemic areas ( $p = 0.00$ )

**Table 5:**
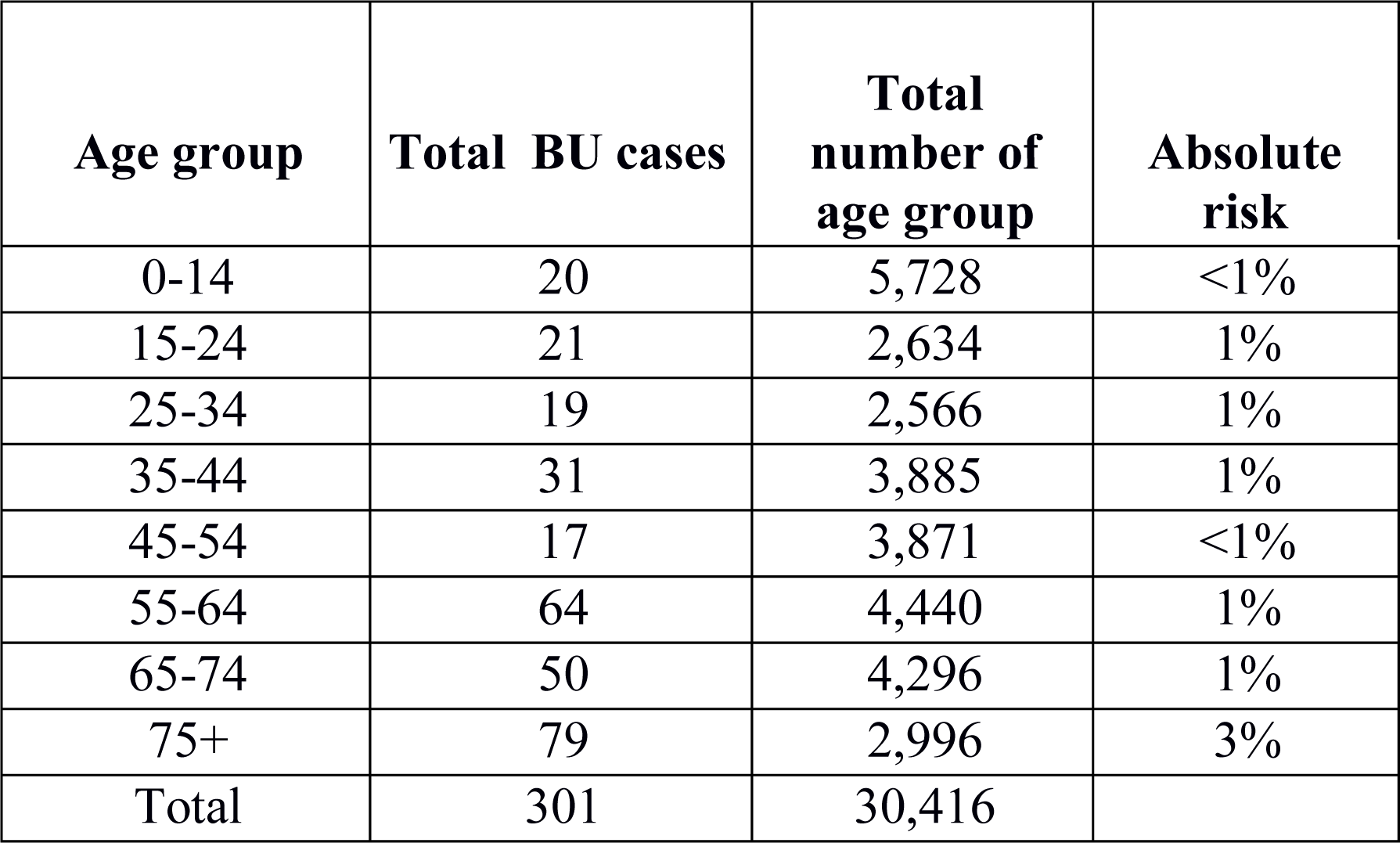
Buruli ulcer absolute risk for age group for the top five *Residents’* location combined using 2016 population census data.

**Table 6:**
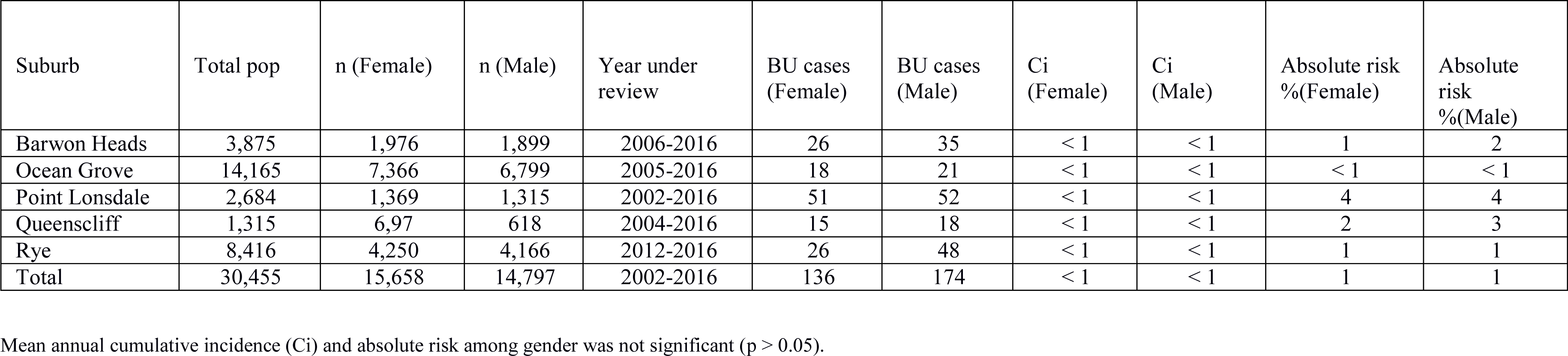
Buruli ulcer incidence rate and absolute risks for gender in the top five *Residents’* locations.

| Suburb | Total pop | n (Female) | n (Male) | Year under review | BU cases (Female) | BU cases (Male) | Ci (Female) | Ci (Male) | Absolute risk %(Female) | Absolute risk %(Male) |
| --- | --- | --- | --- | --- | --- | --- | --- | --- | --- | --- |
| Barwon Heads | 3,875 | 1,976 | 1,899 | 2006-2016 | 26 | 35 | < 1 | < 1 | 1 | 2 |
| Ocean Grove | 14,165 | 7,366 | 6,799 | 2005-2016 | 18 | 21 | < 1 | < 1 | < 1 | < 1 |
| Point Lonsdale | 2,684 | 1,369 | 1,315 | 2002-2016 | 51 | 52 | < 1 | < 1 | 4 | 4 |
| Queenscliff | 1,315 | 6,97 | 618 | 2004-2016 | 15 | 18 | < 1 | < 1 | 2 | 3 |
| Rye | 8,416 | 4,250 | 4,166 | 2012-2016 | 26 | 48 | < 1 | < 1 | 1 | 1 |
| Total | 30,455 | 15,658 | 14,797 | 2002-2016 | 136 | 174 | < 1 | < 1 | 1 | 1 |

Mean annual cumulative incidence (Ci) and absolute risk among gender was not significant ( $p > 0.05$ ).

## DISCUSSION

### Progression of disease in Victoria

Buruli ulcer continues to be reported in Victoria after the first cases in Bairnsdale area of East Gippsland in the 1930s (13,14). Previously studies have concentrated in focal areas, especially in the Bellarine Peninsula and more specifically Point Lonsdale to understand the progression and transmission of the disease (12,19,32). The incidence rate in Victoria from 1994 to 2016 was 0.8/100,000 persons. Cases in Point Lonsdale have been decreasing and there is a gradual movement of BU case towards south-eastern Victoria and the reason is unclear. Though the BU database provides disease cases from 1994, temporal analysis identified that disease clustering became significant in 2011 (**Fig 4**). However space-time analysis reports disease clustering in Bellarine and Mornington Peninsulas from 2001 to 2016 (**Fig 3**). Some of the secondary clusters detected may indicate *Residents’* not living in known endemic areas with exposure to new endemic regions (**Fig 3**). It is worth noting that increase in cases has not been uniform in Victoria and there are top five endemic locations (**Fig 5**). In addition, the most likely disease cluster detected circumscribed the Bellarine and Mornington Peninsulas (**Fig 3**). The reason for such a shift is unclear. It is plausible that the high concentration of people in metropolitan areas, increase in population and construction of new settlements in these areas could play a role to recent rise in cases. This is evident from the demographic changes that have taken place since 2006. There was no direct correlation between incidence rate and rainfall data analysed at three data points at the local scale presented in this study. We are aware that, to expect incidence rate to rise with rainfall pattern alone, is an oversimplification as the transmission pattern of *M. ulcerans* to humans could be far more complex and may involve several critical points at larger scale (33) and intermediate hosts as described by Roche *et al.,* 2013 (34).

The month with the highest notifications in the BU database was August. Previous studies have reported the median incubation period of BU to be 4.5 months (range: 1 to 9 months) after infection (35). This suggests that more people would have been exposed or infected in autumn, March-April (4.5 months median lag period) or November-February (9 months maximum lag period), which is the summer period in Victoria. This was similar to previous studies (15,35,36). This is not surprising as more outdoor activities occur during the summer and autumn periods. Increased contact with the environment was observed from the activities undertaken by *Visitors* in the perceived exposure sites were mostly holiday related (data not shown). Comparison of the month of notification among *Residents* and *Visitors* for the top 5 exposure sites (**Fig 6**) showed close similarity to the trend in Victoria (**Fig 2D**). This may confirm that exposure sites of *M. ulcerans* is likely to be in the top five *Residents’* locations (**Fig 5A**).

However, **Fig 2D** may suggests that as cases increases in Victoria, more new suburbs are detected among (*Residents*). This trend could be interesting and may suggest wider foci of risk factors across Victoria than probably suspected. These new suburbs may describe breeding grounds for the next outbreak.

### Age and gender disparities in BU cases in Victoria

Though males reported more cases within the database, there was no statistically significant difference in incidence rate and absolute risk between genders. A closer look at *Residents’* and *Visitors’* profile of the top five endemic areas for gender, showed no significant difference as well (**Table 2**) indicating that gender did not appear to be a risk factor for BU. Age was a risk factor in the endemic areas (**Table 4**, p = 0.00). The association of age and gender in BU transmission have previously been reported both in Africa and Australia (22,37). In Africa, gender was not a risk factor however, age was a risk factor especially among the less than 15 and over 79 year age groups (6,22). Previous studies in Victoria also showed no difference in gender, but the over 60 age group were most affected. In Australia, this has been attributed to immunosenescence (37,38). Our findings agree with previous studies in Victoria. The findings in this study limits generalization of BU case reports in Victoria, because the trend observed in the State is heavily influenced by only a few suburbs and highly focal. Absolute risk was unusually high (4%) among the 15-24 age group in Point Lonsdale which has never been reported before.

## CONCLUSION

We acknowledge that BU incidence rate from 1994 to 2004 will be underestimated as in these years we relied on voluntary reporting (this would have missed a significant number of cases). After the identification of the first BU cases 86 years ago, BU still presents a major public health concern. There has been a dramatic increase in number of BU cases since 2010. The debilitating features and associated complication presents a significant burden to patients and subsequently high medical costs (39,40). This study has provided a detailed profile of the epidemiology of BU cases in Victoria utilizing the BU database (1994 to 2016). Increasing population and construction of new settlements may have contributed to the recent BU rise in some areas. Gender is not a risk factor, however age is. The age group most affected in Victoria were the ≥ 60 years. This has been attributed to immunosenescence. The distribution pattern of BU cases or emergence observed in our study is not random, as have previously been reported (33). Spatial disease clusters have been detected in Victoria and for the first time they have been tested and four spatial clusters have been found significant. The most likely cluster is on the Bellarine and Mornington Peninsulas.

The incidence rate observed in the most likely disease cluster area was 40/100,000 persons from 1994 to 2016 (space-time analyses). The relative risk for the top five locations combined was 101. There is a strong confirmation of the top five patients’ location (*Residents*) being the environmental source of *M. ulcerans* exposure. The identification of specific environmental factors is still lacking. It is plausible that behavioural patterns of *Residents* and *Visitors* could provide key information of specific risk factors. Analysing the daily activities or lifestyle pattern of *Residents* requires further study. Therefore, future case control studies could be designed to describe the daily activities of *Visitors* (e.g. hiking or swimming, where and when) during their stay. This could provide valuable information regarding identification of possible risk factors. In addition, since *M. ulcerans* is an environmental mycobacteria, extensive environmental sampling would be necessary to determine the exact source of transmission to humans for public health intervention.

## Acknowledgements

Our profound appreciation to the Department of Health & Human Services, the Victorian *Mycobacterium* Reference Laboratory, notifying doctors and primary laboratories for generating the notification, demographic and risk factor data used in this study. Our profound appreciation to Melanie Thomson for her involvement in securing the BU notification database for the study. We are grateful to Fiona Collier and StellaMay Gwini (Barwon Health, Geelong, Australia), Anthony Chamings (Geelong Centre for Emerging Infectious Diseases, Geelong, Australia) and two other anonymous individuals for their insightful comments and suggestions on the descriptive epidemiology analyses for their insightful comments and suggestions on the descriptive epidemiology analyses.

## Supporting information

S1 Checklist. STROBE checklist.

